# Multi-omics approaches define novel aphid effector candidates associated with virulence and avirulence phenotypes

**DOI:** 10.1101/2024.07.30.605808

**Authors:** Peter Thorpe, Simone Altmann, Rosa Lopez-Cobollo, Nadine Douglas, Javaid Iqbal, Sadia Kanvil, Jean-Christophe Simon, James C. Carolan, Jorunn Bos, Colin Turnbull

## Abstract

**Background:** Compatibility between plant parasites and their hosts is genetically determined by both interacting organisms. For example, plants may carry resistance (R) genes or deploy chemical defences. Aphid saliva contains many proteins that are secreted into host tissues. Subsets of these proteins are predicted to act as effectors, either subverting or triggering host immunity. However, associating particular effectors with virulence or avirulence outcomes presents challenges due to the combinatorial complexity. Here we use defined aphid and host genetics to test for co-segregation of expressed aphid transcripts and proteins with virulent or avirulent phenotypes.

**Results:** We compared virulent and avirulent pea aphid parental genotypes, and their bulk segregant F1 progeny on *Medicago truncatula* genotypes carrying or lacking the *RAP1* resistance quantitative trait locus. Differential gene expression analysis of whole body and head samples, in combination with proteomics of saliva and salivary glands, enabled us to pinpoint proteins associated with virulence/avirulence phenotypes. There was relatively little impact of host genotype, whereas large numbers of transcripts and proteins were differentially expressed between parental aphids, likely a reflection of their classification as divergent biotypes within the pea aphid species complex. Many fewer transcripts intersected with the equivalent differential expression patterns in the bulked F1 progeny, providing an effective filter for removing genomic background effects. Overall, there were more upregulated genes detected in the F1 avirulent dataset compared with the virulent one. Some genes were differentially expressed both in the transcriptome and in the proteome datasets, with aminopeptidase N proteins being the most frequent differentially expressed family. In addition, a substantial proportion (27%) of salivary proteins lack annotations, suggesting that many novel functions remain to be discovered.

**Conclusions:** Especially when combined with tightly controlled genetics of both insect and host, multi-omics approaches are powerful tools for revealing and filtering candidate lists down to plausible genes for further functional analysis as putative aphid effectors.

## Background

Crop losses due to insect pests represent an enduring challenge for agriculture and global food security. Aphids are a major problematic group, due both to the direct damage they cause by phloem sap feeding and to indirect effects through acting as vectors for transmission of many viruses. Impacts of pests are further exacerbated by the breakdown of genetically based crop resistance mechanisms due to selection pressures driving pest evolution, as well as evolved insecticide resistance.

In contrast to related fields such as plant-pathogen interactions, the molecular relationships that determine (in)compatibility of plant-aphid interactions are relatively poorly understood. Specific resistance to plant pathogens frequently involves recognition of pathogen effectors, often by resistance proteins (R) characterised by nucleotide-binding and leucine rich repeat (NLR) domains. Several coiled coil domain NLR proteins have been implicated in resistance to aphids and their close relatives. For example, Mi-1, Vat and Bph14 confer resistance to certain biotypes of *Macrosiphum euphorbiae* (potato aphid) [1], *Aphis gossypii* (melon-cotton aphid) [2] and *Nilaparvata lugens* (brown planthopper) [3], respectively. These NLR receptors are predicted to be involved in direct or indirect recognition of molecular signatures that insects, like plant pathogens, release inside their hosts. Indeed, aphids secrete multiple effector proteins into their saliva, that are then predicted to be delivered into plant tissues to modulate host cell processes and to suppress or trigger host defences [4–7]. Although there is one recent report of the BISP effector from brown planthopper, an aphid relative, interacting with the BPH14 NLR in rice [8], there are currently no examples where cognate aphid effector and NLR pairs have been fully defined. Improved molecular insights into virulence and resistance mechanisms taking place during both compatible and incompatible plant-aphid interactions are therefore a priority, and can provide essential knowledge for future development of durable aphid control strategies.

The availability of extensive genome, transcriptome and resequencing resources for the model aphid species *Acyrthosiphon pisum* (pea aphid) [9, 10] have enabled comprehensive genome-wide explorations. There are also genomic sequences now available at NCBI and AphidBase (https://bipaa.genouest.org/is/aphidbase/) for more than 25 species of aphids and close relatives, often associated with gene predictions and transcriptomes [11]. In addition, several papers have attempted to define the aphid effectorome, either by direct analysis of salivary proteins, or by transcriptomics of salivary glands, coupled with filters for predicted secreted, non-trans-membrane proteins [12–17]. Beyond the true aphids (superfamily Aphidoidea), there are now genomic resources for sister groups within the Hemiptera such as planthoppers, leafhoppers, psyllids, whitefly and scale insects (https://www.ncbi.nlm.nih.gov/assembly/?term=hemiptera) that likewise are major crop pests, alongside genomes for triatomines and bed bugs, hemipterans that feed on animal rather than plant hosts. Outside the Hemiptera, genomic data have been published for sucking pests such as thrips and spider mites that feed on plant tissues other than phloem [18–20]. Genome, transcriptome and proteome comparisons across clades may enable definition of putative effector subsets that are necessary for different feeding modes, and may provide insights into conserved and divergent modes of action in terms of how the plant immune system is targeted to enable successful parasitism.

Despite the wide range of functional genomics studies published to date, one common limitation is the lack of understanding of the differences in effector complements between virulent (host-compatible) and avirulent (host-incompatible) genotypes. Genetic differences operate at several taxonomic levels. First, there are major differences across aphid species in their host preferences and host compatibilities. Some species, such as peach potato aphid (*Myzus persicae*) are generalists that can feed on at least 400 known plant species, making them widespread crop pests [21]. Others are specialists, such as pea aphid (*A. pisum*) that exclusively feeds on legumes (Fabaceae). Second, there is substantial diversity within species such as *A. pisum* that has led to its description as a species complex comprising several host races that each have a strong preference for particular legume species, supported by robust molecular marker fingerprints for each host race [22, 23]. There is evidence of divergence and differential expression of chemosensory gene families such as odorant receptors across different pea aphid biotypes [24, 25], but causative relationships have yet to be established for genes and proteins that govern the range of compatible and incompatible interactions seen. There is also clear evidence that some host races can survive and sometimes thrive as migrants on hosts outside their preferred species range [22]. Finally, at the intra-specific level for both aphids and hosts, there can be a wide range of compatibilities. For example, from testing eight genotypes of *A. pisum* in combination with 23 different *Medicago truncatula* (Mt) accessions, we discovered high diversity in both species that did not correspond particularly strongly to host races or to geographic origins of the host lines [26]. Parallel to this, crossing two divergent pea aphid biotypes to generate F1 recombinant populations uncovered Mendelian segregation of virulence/avirulence on Mt genotypes carrying the *RAP1* aphid resistance QTL [27, 28].

Here, we report global exploration of the molecular basis for aphid virulence and avirulence on defined host genotypes. Specifically, we aimed to link phenotypes to candidate effectors and related genes by multiple comparisons of the transcriptomes and proteomes of two divergent parental pea aphid clones, along with the transcriptomes of segregating avirulent and virulent pooled individuals from within F1 cross populations (Fig. 1). We also critically analysed the effectiveness of combined omics approaches as a means to robustly uncover proteins with pivotal biological roles, such as effectors that determine the difference between virulent and avirulent outcomes.

**Figure 1.**
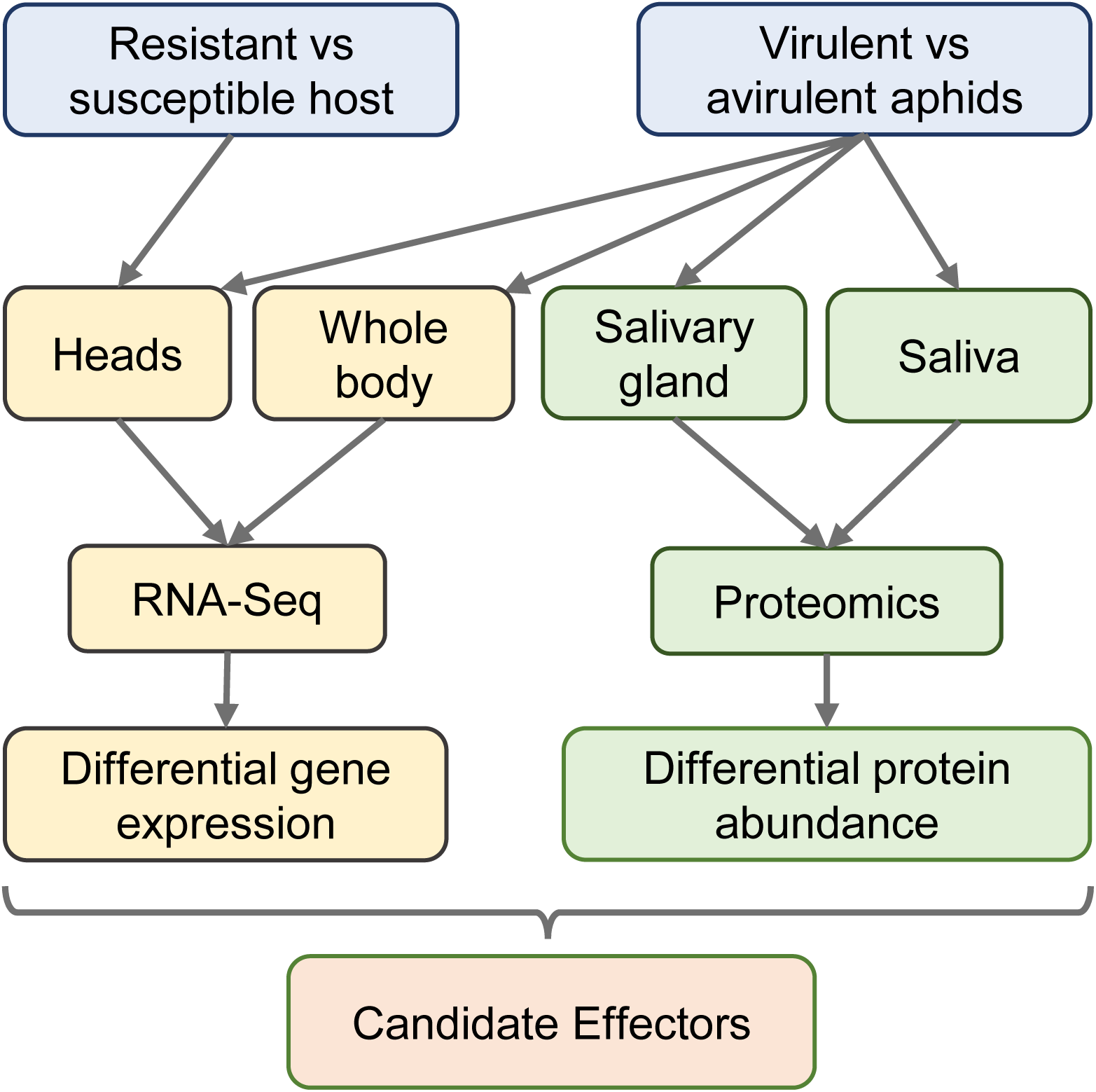
Summary of transcriptome and proteome analysis pipeline. Resistant and susceptible host plants carried or lacked the *RAP1* aphid resistance QTL, respectively. For all experiments, virulent N116 and avirulent PS01 aphids were compared. In addition, BSA-RNA-Seq was done on whole body pooled samples of F1 virulent and avirulent aphids.

## Results and Discussion

### Generation and analysis of aphid populations for RNA-Seq analyses

In our previous work [27], we had demonstrated Mendelian segregation of inheritance of virulent and avirulent phenotypes in F1 pea aphid populations derived from a cross between N116 and PS01 (virulent and avirulent parental clones, respectively) when infested on *M. truncatula* hosts carrying the *RAP1* resistance QTL [28]. On this basis, we reasoned that the molecular basis of the difference between virulent and avirulent aphids could be revealed by transcriptomic and proteomic analysis. However, there were likely to be thousands of genetic and gene expression differences between the parental genotypes, that are representatives of phenotypically contrasting biotypes within the highly diverse pea aphid species complex [22, 26]. This makes it difficult to discern unrelated genomic background differences from causative genes responsible for suppressing host immunity or for triggering R-gene dependent defences. To address this challenge, we employed a bulk segregant analysis (BSA-) RNA-Seq approach that would both reduce the genetic background effects and allow us to test for heritability of differentially expressed (DE) genes across parental and F1 generations. Enabling this strategy first required us to re-create the segregating F1 populations previously reported [27].

We induced sexual forms of PS01 and N116 and conducted reciprocal crosses, leading to screening of a total of 78 F1 clones on two host plant genotypes carrying *RAP1*: Jemalong A17 (hereafter A17), the original source of the identified *RAP1* QTL, and a resistant near-isogenic line (RNIL) derived from a mapping population [29] using A17 as one of the parents. The *RAP1* aphid resistance QTL is highly effective against PS01 aphids, typically resulting in high mortality, whereas N116 aphids are unaffected. Progeny were verified as true F1 hybrids by a panel of seven SSR markers [22] and by screening for maternal inheritance of secondary symbionts reported in the pea aphid [30]. Using a virulence index based on a combination of aphid survival and reproduction, F1 clones were first ranked according to performance on A17. Phenotypes ranged from fully virulent to fully avirulent (Supplementary Material 1A), similar to previous findings [27], although in the present experiment the population as a whole did not display complete segregation into discrete virulent and avirulent categories. As also previously shown, resistance in the RNIL was slightly weaker than in A17, with F1 clones ranging from virulent to avirulent, and importantly performance on the two host genotypes was significantly correlated (Pearson r 0.72, P 1.82e^-13^). All F1 clones were virulent on hosts lacking *RAP1* (Supplementary Material 1B). We then selected 22 sibling F1 clones from each end of the distribution to provide two bulk sample sets with the strongest virulent (VIR) and avirulent (AVR) phenotypes for subsequent transcriptomic analysis. Fig. 2 shows the complete separation of the selected clones into virulent and avirulent classifications. As a final check prior to RNA-Seq experiments, we re-confirmed separation of survival rates of these two subsets of clones on both resistant host genotypes (Supplementary Material 1C).

**Figure 2.**
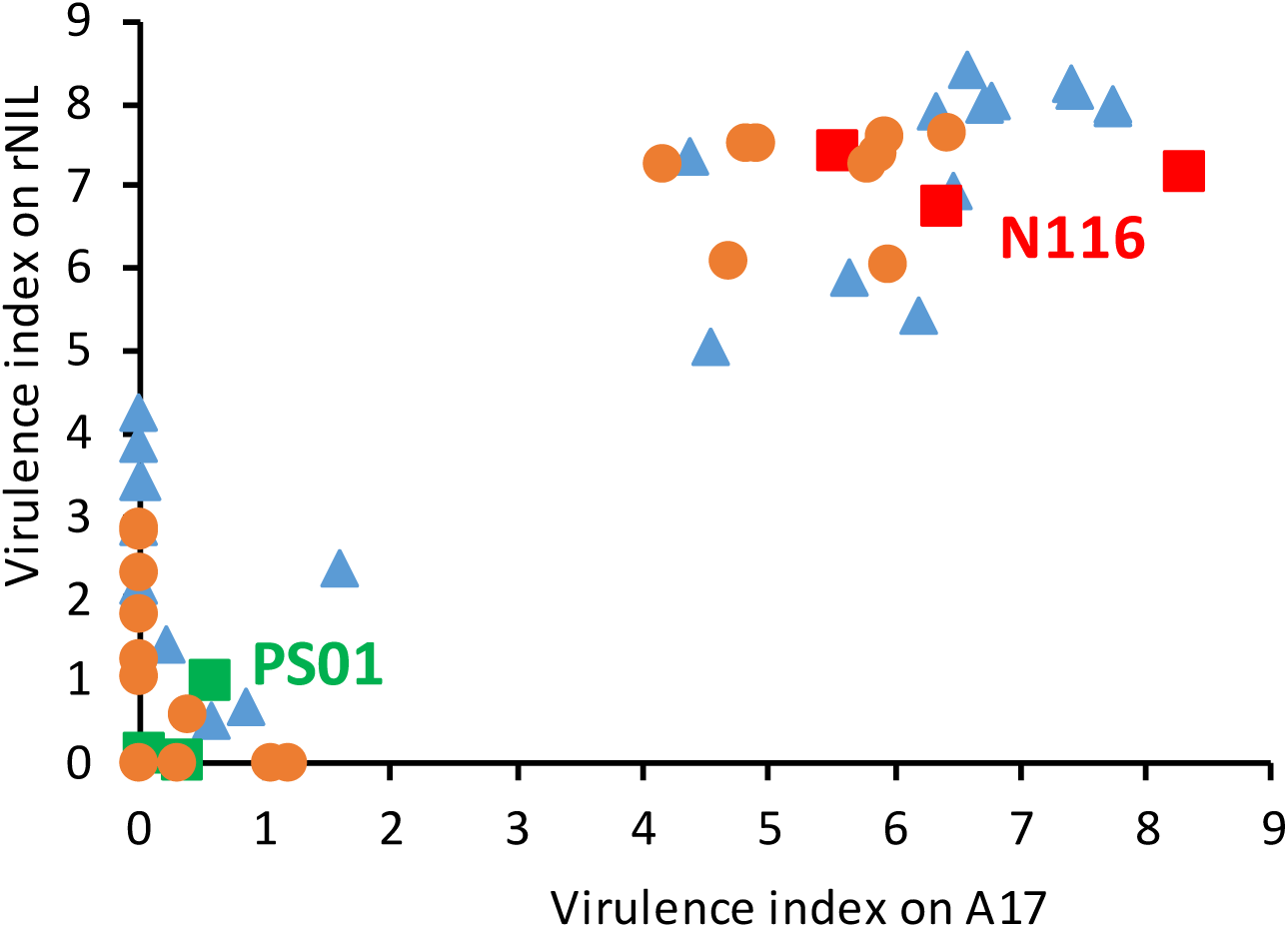
Virulence phenotypes of parental aphid clones and selections from F1 populations used for BSA-RNA-Seq. Tested on two *M. truncatula* genotypes carrying the *RAP1* locus: Jemalong A17 and a resistant near isogenic line (RNIL) derived from a cross between A17 and DZA315.16. The parental genotypes and selections from the F1 populations shown here were all used for the BSA-RNA-Seq experiment. Data are expressed as virulence index, assessed 10 d after infestation. Phenotypes of F1 clones were classified using the following virulence index cut-offs: A17 VIR >4, AVR <2; RNIL VIR >4, AVR <4.5. Orange circles are NP (N116 female x PS01 male); blue triangles are PN (PS01 female x N116 male); red is N116, and green is PS01, with each of three parental data points from a separate batch of F1 tests. The full population phenotype data are provided in Supplementary Material 1.

### Transcriptomic analyses

We first ran an RNA-Seq experiment using the parental clones N116 and PS01 infested onto either A17 or the susceptible DZA315.16 host (hereafter DZA) for 24 h prior to collection of heads for RNA extraction. The multiple aims were to enrich for transcripts from salivary glands that express candidate effectors, to uncover the transcriptome differences between the parental aphid genotypes, and to reveal the impact of host plant genotype. Each aphid x host combination was replicated three times, giving a total of 12 libraries, ranging from 6.8 to 10.6 million reads uniquely mapped to the reference genome (Supplementary Material 2A).

Hierarchical clustering and principal components analysis (PCA) of the transcriptomic expression profiles both indicated that the replicates of each treatment were closely correlated in all cases, so no datasets needed to be discarded (Fig. 3A,B). These analyses additionally revealed that samples were separated largely by aphid genotype rather than host plant treatment. Overall, the transcriptomes of the two aphid genotypes on A17 plants were clearly differentiated, with a total of 483 genes significantly upregulated in N116 and 452 in PS01 (log2 fold change >2.0, FDR <0.05; Supplementary Material 3; Fig. 3C). Similarly, on DZA host plants, 395 and 363 genes were upregulated in N116 and PS01, respectively. In contrast, expression of relatively few genes, between three and 27, across all the pairwise comparisons, was significantly affected by the host plant (Supplementary Material 3; Fig. 3C). Functions of the DE genes are considered below, in conjunction with the other transcriptomic and proteomic experiments.

**Figure 3.**
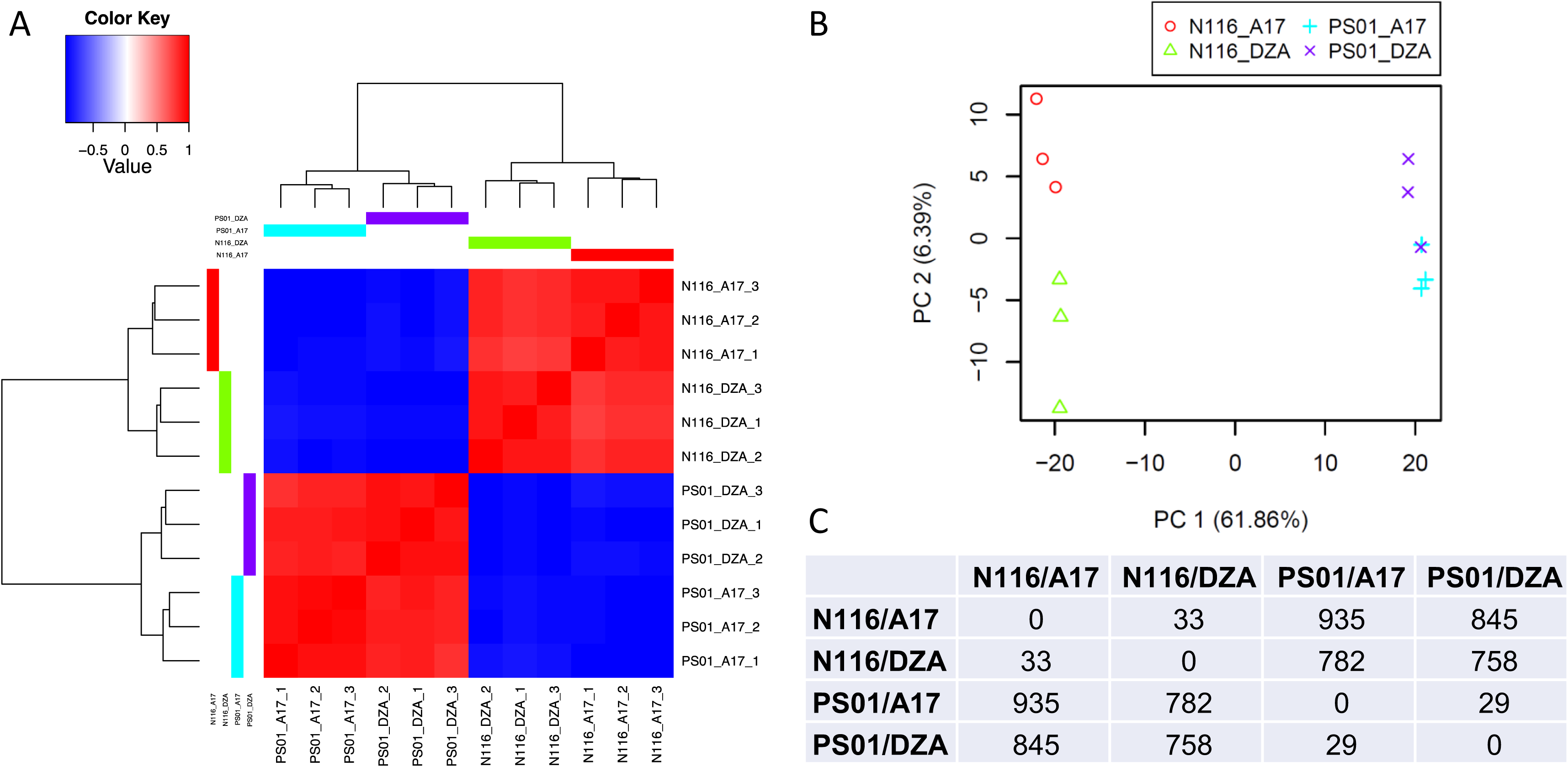
Transcriptome analysis of aphid heads. Samples were dissected heads from PS01 and N116 genotypes infested on *Medicago truncatula* A17 or DZA315.16 for 24 h, with n=3 biological replicates. Aphid genotype PS01 is avirulent on *M. truncatula* A17, and all other combinations represent compatible interactions. A. Clustering of transcriptional responses of pea aphid, showing samples clustered more strongly based on aphid genotype than on host interaction; B. Principal components analysis. The top two principal components explain >68% of the variation among transcriptional responses. Samples group largely by aphid genotype rather than host interaction; C. Numbers of genes differentially expressed between the different aphid genotypes on different *M. truncatula* genotypes. Of the 935 DE genes between PS01 and N116 on A17, 483 were up in N116 and 452 were up in PS01. Of the 758 DE genes on DZA hosts, 395 were up in N116 and 363 were up in PS01. Accompanying gene lists and annotations are provided in Supplementary Material 3.

We next undertook a larger RNA-Seq experiment, sampling whole aphid bodies in order to capture transcripts from all tissues. Using aphids infested onto A17 host plants for 24 h, we again compared N116 and PS01 parental clones, but this time alongside the bulked segregant pools of VIR and AVR F1 clones described above. Five biological replicates for each gave a total of 20 RNA libraries each containing 14 to 22 million reads that uniquely map to the reference genome (Supplementary Material 2B).

Similar to the heads experiment, multivariate analysis by hierarchical clustering and PCA both indicated that all replicates within each sample type grouped together, and that each sample type was clearly differentiated. As expected, the genetically divergent parents were again highly separated, whereas the two pooled F1 datasets were much closer to each other, as they contain 50% of each parental genome, with each pool representing the average transcriptome of multiple independent F1 clones (Fig. 4A,B).

**Figure 4.**
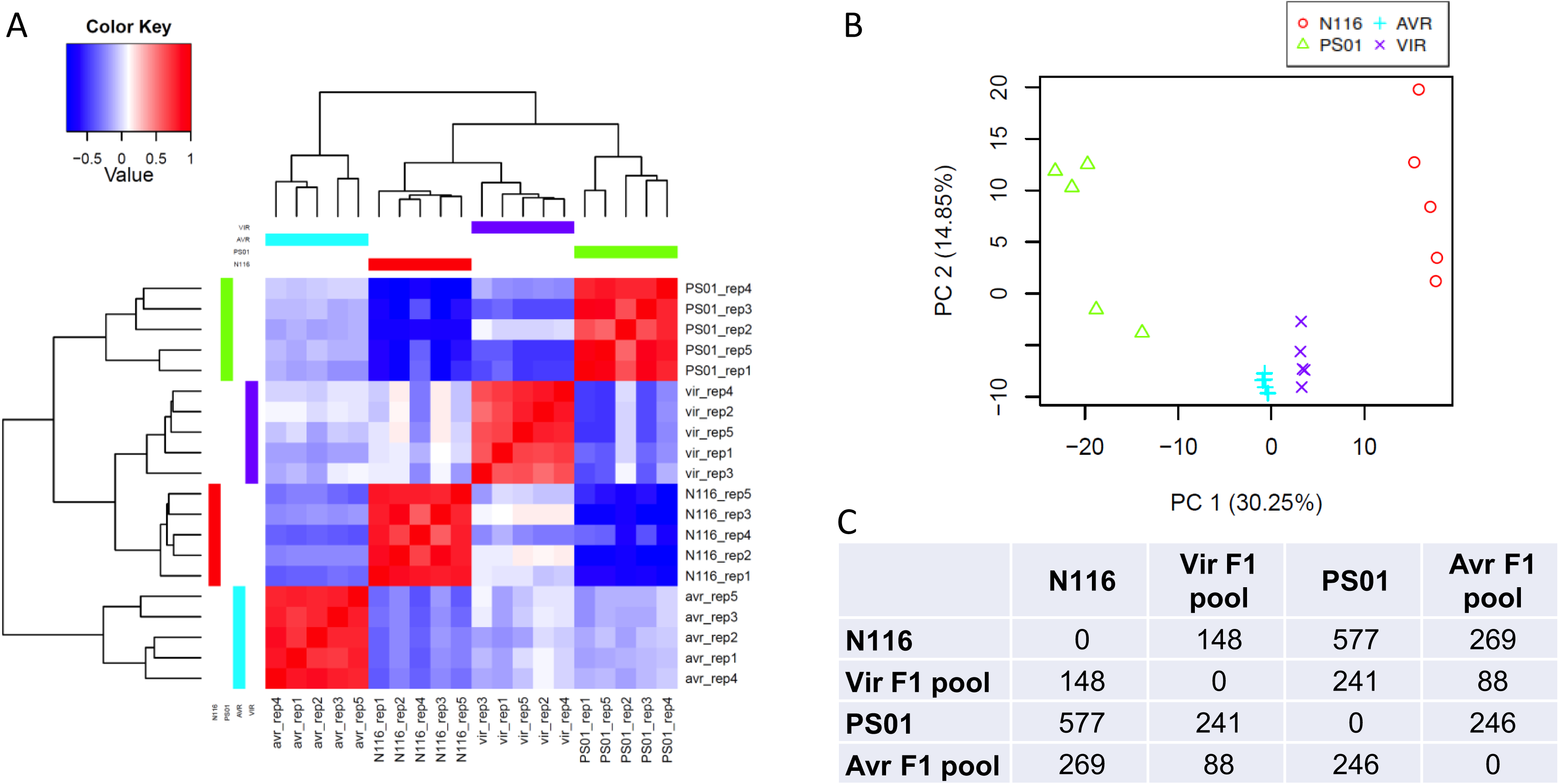
Transcriptome analysis of whole aphids. Aphids were infested on *Medicago truncatula* A17 for 24 h, with n=5 biological replicates. A. Clustering of transcriptional responses of pea aphid PS01, N116, bulked F1 VIR and AVR progeny. Responses within biological replicates are more strongly correlated than responses among different aphid genotypes; B. Principal components analysis. The top 2 principal components explain >45% of the variation among transcriptional responses of PS01, N116, AVR and VIR F1 progeny replicates, and separate the responses of the different aphid genotypes and F1 pools. C. Numbers of genes differentially expressed between the different aphid genotypes and pools. Accompanying gene lists and annotations are provided in Supplementary Material 3.

Differentially expressed genes were identified for all pairwise comparisons between samples (Fig. 4C). The number of up and down-regulated genes between the parental pairs and the pair of F1 pools are shown in Fig. 5A, with the gene lists provided in Supplementary Material 3. Several hundred genes were differentially expressed in both the whole-body and head comparisons of the parents. Some of these DE genes likely reflect genomic differences between the parental clones that are representatives of divergent pea aphid biotypes. However, relatively few DE genes were detected in the F1 samples, with only 24 genes up-regulated in the VIR pool and 64 in the AVR pool. These numbers can also be interpreted as a higher number of genes being down-regulated in the VIR F1 aphids. Fig. 5B,C show the overlaps across head and whole-body datasets for N116/VIR and PS01/AVR, respectively. Unexpectedly, the intersections of DE genes revealed subsets where the direction of expression was opposite between the parental pair and the F1 pooled pairs, with three genes upregulated in N116 and AVR F1, and 13 genes upregulated in PS01 and VIR F1 (Fig. 5D, Fig. 7G,H). Moreover, very few genes were upregulated in both parental N116 and VIR F1 pool datasets. A plausible explanation is that the genes governing virulence in N116 are not the same as those that result in virulent phenotypes in the F1 population. Each individual in the F1 population carries a random 50% of the genome of each parent, creating a high degree of combinatorial complexity. Nonetheless, the DE genes in the F1 data derive from the average across the 22 individuals used to create each bulk RNA pool, and are therefore likely to be biologically relevant to virulence or avirulence functions rather than background genomic noise. Such genes merit further exploration in both parental and F1 genotypes.

**Figure 5.**
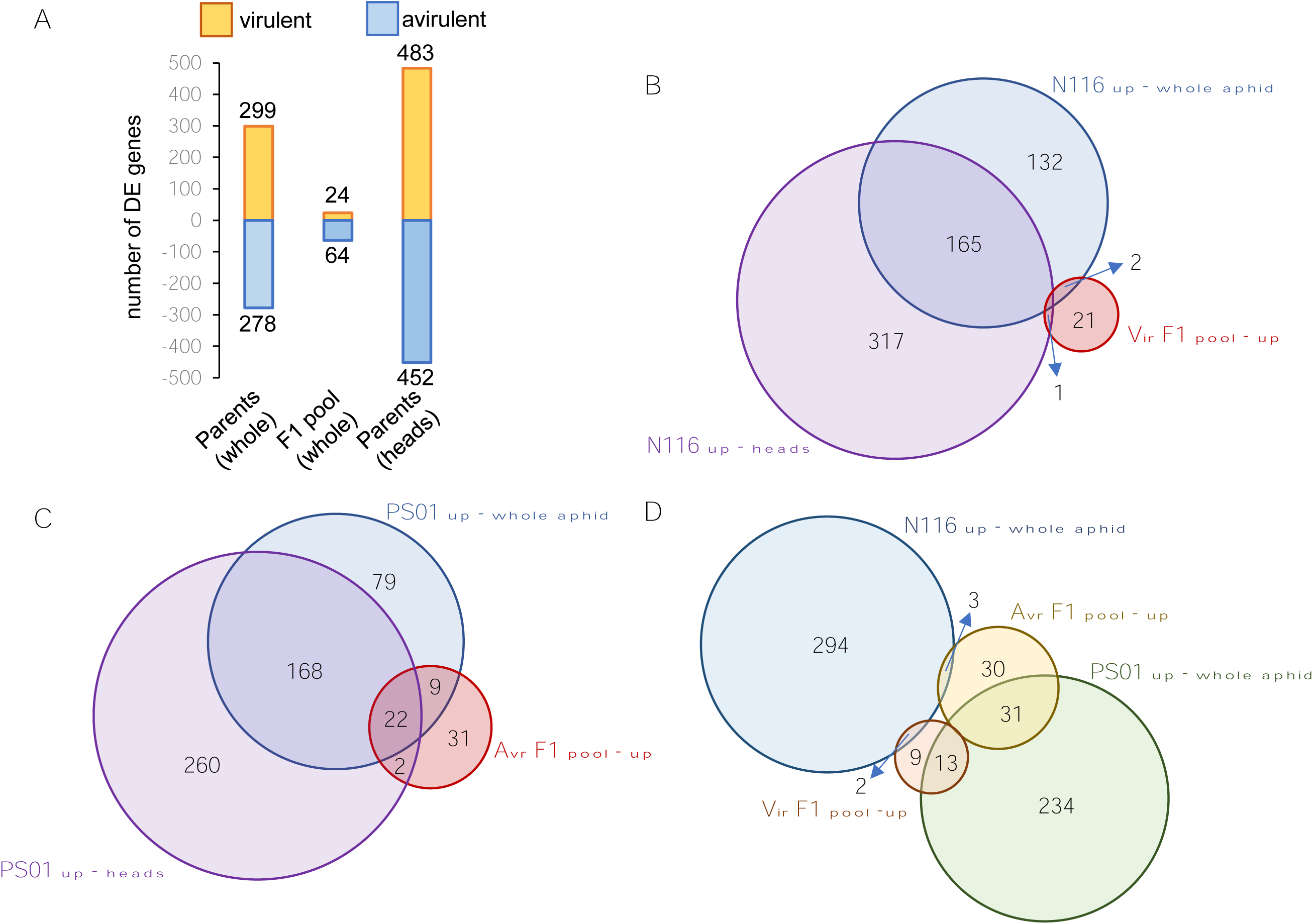
Differential gene expression in pea aphid genotypes N116, PS01 and bulked F1 pools of virulent and avirulent progeny. A. Numbers of genes up-versus down-regulated in comparisons of parent genotypes N116 (virulent) and PS01 (avirulent), and their F1 progeny pools, all on A17 host plants. Orange bars represent the numbers of genes up-regulated in genotype N116 or the VIR F1 pool compared with the genotype PS01 and the AVR pool, respectively. Blue bars represent the numbers of genes up-regulated in genotype PS01 or the AVR F1 pool compared with the genotype N116 and the VIR pool, respectively; B. Overlaps in genes up-regulated in genotype N116 whole body and head tissues compared to genotype PS01, and up-regulated in the VIR F1 pool compared to the AVR F1 pool; C. Overlaps in genes up-regulated in genotype PS01 whole body and head tissues compared to genotype N116, and up-regulated in the AVR F1 pool compared to the VIR F1 pool; D. Overlaps in up-regulated genes among whole body transcriptomes of N116, PS01, VIR F1 pool and AVR F1 pool.

### Quantitative proteomic analysis of saliva and salivary glands

To determine whether differences exist between the salivary protein profiles of the two parental aphid clones, a comparative analysis of salivary gland and salivary proteomes was conducted. A total of 2343 and 2276 high confidence proteins were detected from salivary glands of N116 and PS01, respectively (Supplementary Material 4), with 2038 proteins (80%) common to both (Fig. 6A). Each biotype had similar proportions of non-annotated proteins (PS01: 5.4 % and N116: 6.2%) and proteins predicted to have secretion signals (PS01: 16.6% and N116: 17.3%). These proportions of secreted and non-annotated proteins are typical for pea aphid biotypes [12, 31]. Two major clusters were revealed by PCA (Fig. 6C), corresponding to the two aphid genotypes. Principal Components 1 and 2 account for 64% of the variation, indicating distinct protein profiles in the salivary glands of each genotype. This distinction was further supported by quantitative analysis that identified 235 statistically significant differentially abundant (SSDA) proteins (p<0.05), with 136 and 99 proteins having higher abundances in N116 and PS01 salivary glands, respectively (Fig. 6E; Supplementary Material 4). Relative fold changes (RFC) ranged from −48.5 to +140.0 indicating that even when both genotypes engage in compatible interactions with the same plant type (*V. faba* in this case) the salivary gland profiles are divergent both qualitatively and quantitatively.

**Figure 6.**
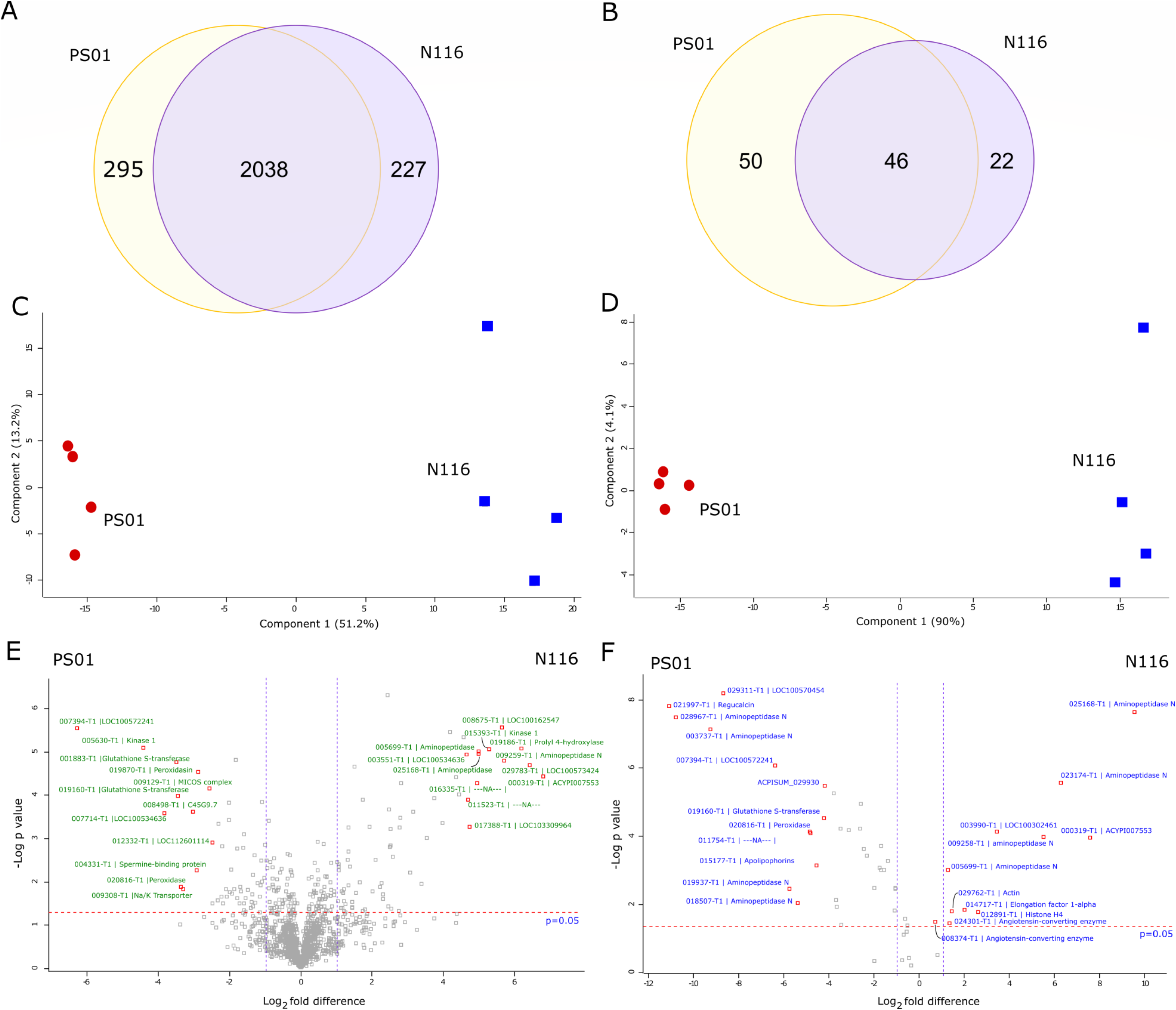
Comparative proteomic analysis of salivary glands and saliva for pea aphid genotypes N116 and PS01. Venn diagrams of the number of proteins shared and found exclusively in A) salivary glands and B) saliva identified for both genotypes. Principal Components Analysis (PCA) of C) salivary glands and D) saliva distinguishes both genotypes clearly. Volcano plots based on -log_10_ p values and log_2_ fold differences highlighting the statistically significant differentially abundant (SSDA) proteins (p≤0.05) for E) salivary glands and F) saliva. Annotations are shown for the top 12 proteins of increased and decreased abundances.

Of the 136 SSDA salivary gland proteins with increased abundance in N116, 60 (44%) were predicted to be secreted and 27 (20%) had no annotations. Similar proportions were observed within the 99 SSDA proteins with increased abundance in PS01, with 33 (33%) and 18 (18%) proteins having a secretion signal or no annotations, respectively. These proportions of secreted and non-annotated proteins within the differentially abundant sets are substantially higher than the corresponding proportions in the background salivary gland proteomes described above. Of the top ten proteins with the highest relative abundance in N116, seven had no annotation: ACPISUM_000319 (ACYPI007553; RFC 140.0) and ACPISUM_029783 (LOC100573424; RFC 64), ACPISUM_008675 (LOC100162547; RFC 32), ACPISUM_016335 (Not annotated; RFC 26), ACPISUM_017388 (LOC103309964; RFC 21.1), ACPISUM_003551 (LOC100534636; RFC, 21.1) and ACPISUM_009099 (LOC112598674, 18.4). The other proteins in the top ten were a kinase ACPISUM_015393 (developmentally-regulated protein kinase 1; RFC 64) and two aminopeptidases (ACPISUM_009259; RFC 36.8 and ACPISUM_005699; RFC 22.6). Of the top ten proteins with highest abundances in PS01 in comparison to N116, two were uncharacterised: ACPISUM_007394 (LOC100572241; RFC 48.5) and ACPISUM_007714 (LOC100534636; RFC 11.3); and two were glutathione S-transferases (ACPISUM_019160 and ACPISUM_001883, both RFCs of 8.6). Other proteins included a different developmentally-regulated protein kinase (ACPISUM_005630; RFC 17.1), a peroxidase (ACPISUM_020816; RFC 9.8), a prostatic spermine-binding protein (ACPISUM_004331; RFC 8), peroxidasin (ACPISUM_019870; RFC 6.5), an ATPase subunit (ACPISUM_009308; RFC 5.7) and glyoxylate reductase (ACPISUM_021751, RFC 4.9). We next examined aphid saliva proteins. Although the samples are collected from artificial diets, these salivary secretomes are likely to be highly similar to the proteins delivered into plant tissues during interactions with the host, and therefore are predicted to include the entire set of effectors. We focussed on categorisation of the total salivary protein lists, and of the DE proteins. Although the analysis of saliva revealed far fewer proteins than from the salivary gland samples, there is again a clear distinction between the two genotypes. A total of 69 and 97 high confidence proteins were found in N116 and PS01 saliva, respectively (Fig. 6B; Supplementary Material 4) with 22 (32% for N116) and 50 (52% for PS01) proteins, being deemed unique to each. A large proportion (30% for PS01 and 25% for N116) of the salivary proteomes had no annotations, indicating their potential phylogenetic restriction to aphids. In addition, 39% and 32% of the proteins had predicted canonical secretion signals for PS01 and N116 saliva, respectively. Notably, although saliva proteins detected in diet samples have, by definition, been secreted, the majority appear not to have canonical secretion signals. Explanations range from incomplete/incorrect gene models to non-canonical or alternative secretion mechanisms. Our results highlight the importance of combining several approaches when attempting to identify potential effectors and molecular determinants of virulence/avirulence. Omitting proteins without secretion signals from bioinformatic pipelines may result in many effector candidates being overlooked.

As with the salivary glands, PCA of the salivary proteins completely resolved two groups, with PC1 and PC2 accounting for 94% of the total variation (Fig. 6D). Label free quantitative analysis using MaxQuant identified 47 SSDA proteins with 12 and 35 proteins having higher abundance in N116 and PS01 saliva, respectively (Fig. 6F; Supplementary Material 4). Notably, N116 saliva comprises fewer detected proteins and fewer SSDA proteins than PS01, possibly pointing to a strategy that enables evasion of host defences. If, for example, one or more of the proteins uniquely detected in PS01 saliva act as avirulence factors due to cognate receptors in the host plant, their absence or low abundance in N116 may result in a compatible interaction. However, it remains to be experimentally determined whether these genotypic differences in type or number of saliva proteins are causatively associated with virulence or avirulence.

Most of the salivary proteins identified here have previously been associated with pea aphid saliva including multiple members of M1 and M2 metalloprotease families, along with peroxidases, glutathione-S-transferases, glucose dehydrogenase and regucalcin [12, 32]. Apart from the Aminopeptidase N (APN) category discussed in detail below, the most frequent annotation was for unknown proteins: 20-26% of the total saliva list for each clone, and 21% of the DE saliva proteins. Four out of the ten DE unknown proteins also featured within the top 20 proteins by MS intensity or protein coverage. High proportions of unknown proteins have been noted in earlier studies of aphid saliva and the salivary gland predicted secretome [31]. In addition, a homologue of a salivary effector previously characterised for *Myzus persicae* (Mp1) [33] had a higher abundance in PS01 saliva (ACPISUM_000421; RFC 14). The relative fold changes of salivary proteins ranged from -2352 for regucalcin to 724 for members of the APN (M1 metalloprotease) family, which represented the most differentially abundant proteins in PS01 and N116 saliva, respectively. Although these RFC values can be considered arbitrary due to imputation of low abundant values in samples where the proteins are in fact absent, there is very clear divergence of salivary proteomes both in the proteins uniquely detected in one or other genotype, and in the large differences in apparent abundance of several proteins present in both genotypes. The full lists of proteins exclusively found in the saliva or salivary gland proteomes of both genotypes are provided in Supplementary Material 4, with 25 and five proteins exclusive to the salivary glands and saliva of N116, respectively. For PS01, the corresponding numbers were 10 and 13 proteins exclusive to the salivary glands and saliva, respectively. These proteins were present in all replicates of one genotype while being absent in all replicates of the other, strongly supporting their status as candidate effectors, that may individually or collectively determine the VIR and AVR phenotypes observed for each genotype on different host plants.

Comparison of the quantitative differences in protein abundance across both the saliva and salivary gland datasets revealed clear similarities in the two proteomes analysed for each genotype. Five proteins that were of higher abundance in N116 saliva were also more abundant in N116 salivary glands in comparison to their PS01 counterparts. A similar trend was observed for nine PS01 salivary and salivary gland proteins (Supplementary Material 4), with the RFCs for these proteins positively correlated across both biological sample types. The fact that the abundances of these salivary gland proteins are mirrored at the level of externally delivered oral secretions highlights the robustness of both analyses, and points to likely roles as virulence or avirulence determinants in two genotypes with distinct host preferences. Such proteins represent excellent candidates for future characterisation to determine their effector status, especially those that are also supported by DE transcript profiles (Table 1).

**Table 1.**
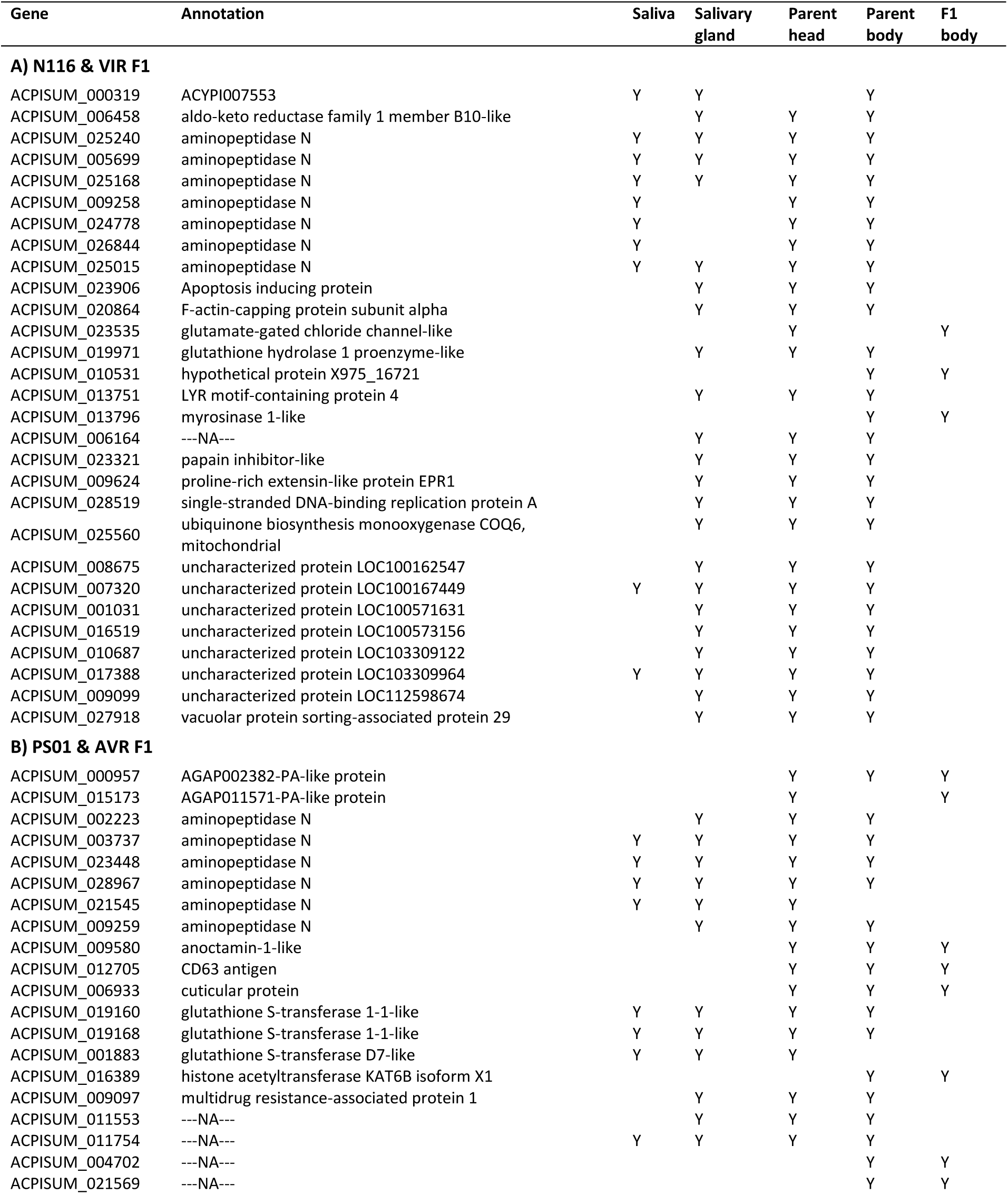
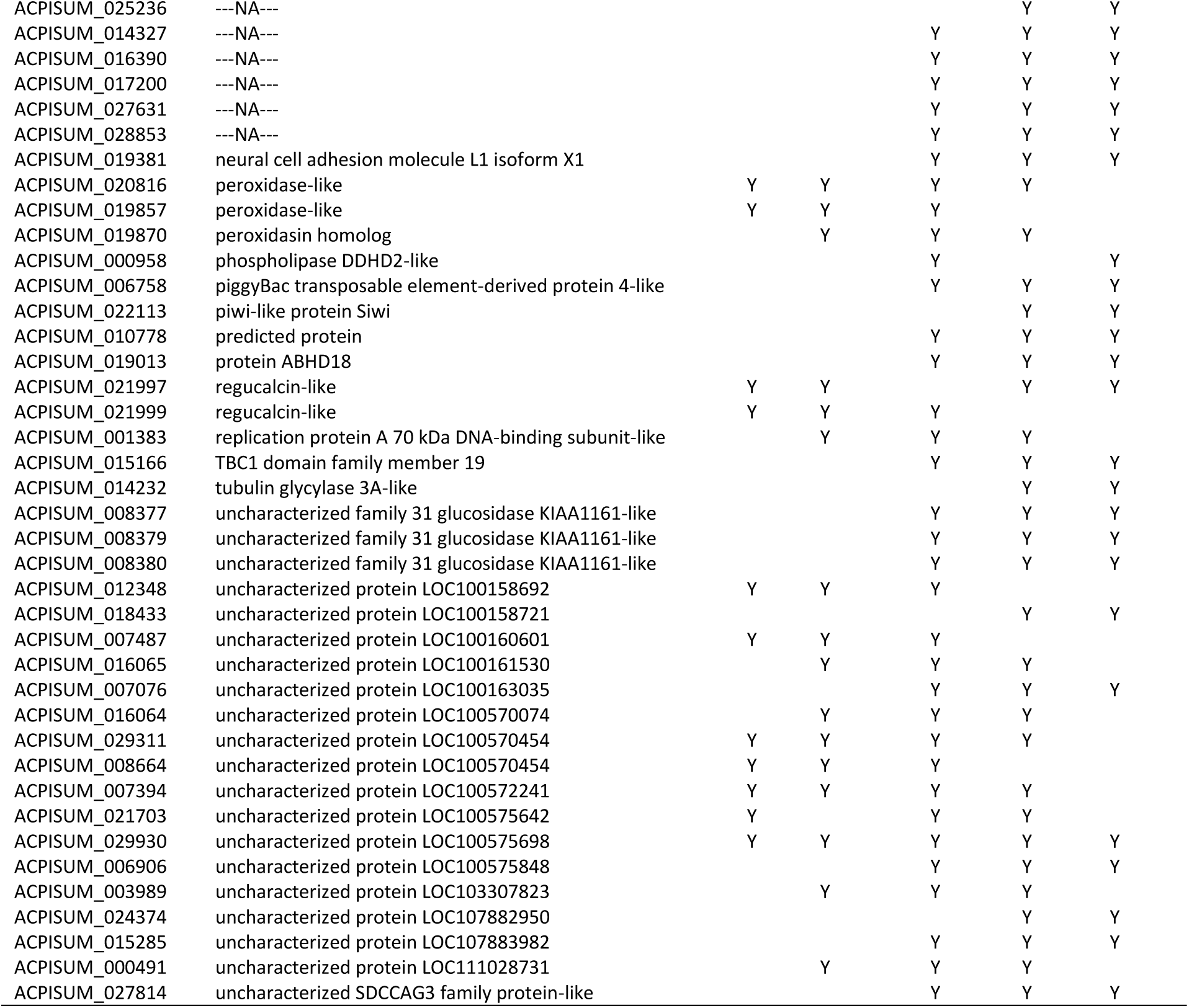
Genes and proteins overlapping in multiple experiments. All genes shown that are represented in at least three datasets, plus all genes intersected between F1 transcriptome and at least one other dataset. Saliva and salivary gland data are proteins, head and body data are transcripts. A. Proteins and upregulated genes in virulent aphids (N116, VIR F1 pool); B. Proteins and upregulated genes in avirulent aphids (PS01, AVR F1 pool). Y = protein present and/or RNA differentially expressed. Full gene and protein lists are in Supplementary Material 3 and 4.

### Overlap between transcriptomics and proteomics datasets

Across the transcriptomics and proteomics experiments, we analysed all the intersections then extracted the proteins and DE gene subsets that showed the greatest overlaps (Table 1; Supplementary Material 3 and 4), partitioning into genes/proteins associated with virulence, in N116 or the VIR F1 pool, or with avirulence, in PS01 or the AVR F1 pool. The number of DE genes or proteins in the head transcriptome, whole body transcriptome and salivary gland proteome datasets were broadly similar between VIR and AVR samples. However, the PS01 saliva protein and the AVR F1 pool transcript lists were longer than those for N116 saliva and VIR F1 pool transcripts, reflected by larger intersections in the former. Over half (33/64) of genes upregulated in the AVR F1 pool were also in at least one other list, whereas only three out of 24 intersected from the VIR F1 pool data. Whole body RNA-Seq data for a selection of these intersected genes are plotted in Fig. 7. Several of the AVR-upregulated genes shown are annotated as enzymes with hydrolase, glycosidase or peroxidase functions. Other annotations include a transcription factor and proteins of unknown function. Genes on the VIR side included ACPISUM_013796 (myrosinase 1-like) and ACPISUM_019971 (glutathione hydrolase 1 proenzyme-like), although these were not found in saliva. Across the multiple experiments, the two most frequently found genes in the AVR data were ACPISUM_021997 (regucalcin-like) previously reported as a Ca-binding protein [32], present in all lists except heads RNA, and ACPISUM_029930 (uncharacterized protein LOC100575698), present in all five lists. These AVR-related salivary proteins represent strong candidates for functional effectors, based on the multiple strands of evidence for their differential expression and importantly for co-segregation of their expression with the avirulence phenotype in the F1 population. We have therefore uncovered heritable differences in salivary proteins that associate with avirulence, in this case an incompatible phenotype on Mt hosts carrying the *RAP1* QTL [27, 28]. Intriguingly, however, we found no equivalent strong candidates for salivary proteins that might represent the dominant virulence factor predicted by previous genetic studies [27]. Alternative explanations for the Mendelian segregation found in that study could be that the proposed “virulence” gene is not an effector per se, but instead could be an upstream positive regulator, or a negative regulator of one or more effectors that act as avirulence factors detected by a *RAP1* dependent pathway.

**Figure 7.**
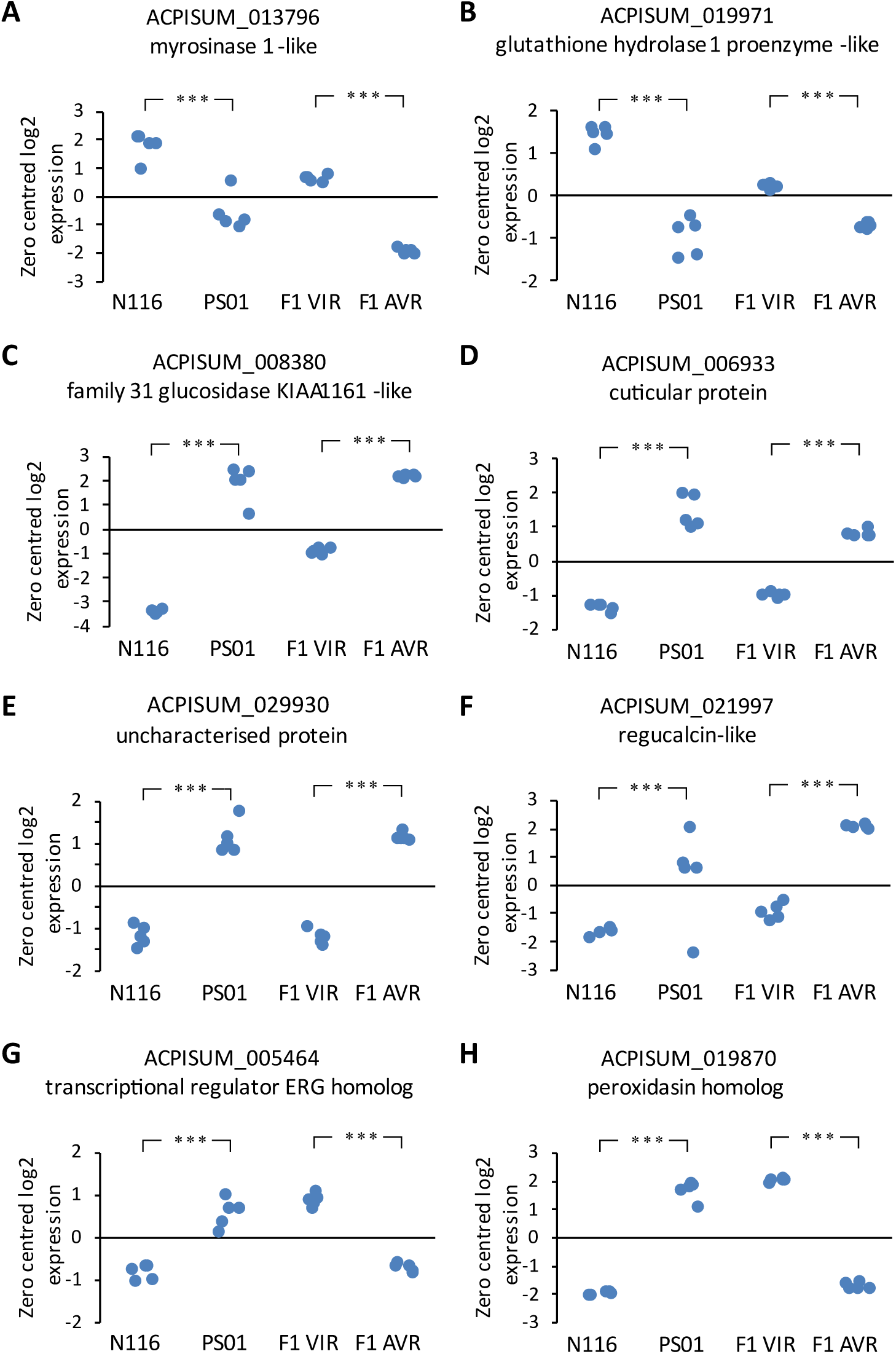
**Selected differentially expressed genes from whole body transcriptomes**. A,B representative genes upregulated both in virulent parent and in virulent F1 pool; C-F representative genes upregulated both in avirulent parent and in avirulent F1 pool. G,H representative genes with opposite regulation between parent and F1 pairs. Each point represents an individual RNA-Seq library (n=5). *** indicates FDR<0.001.

### Gene Ontology analysis

We undertook Gene Ontology (GO) analysis to reveal functional categories and genes that were enriched in the differentially expressed gene and protein data sets. Using a FDR of <0.05, many gene sets contained few or no significantly enriched terms (Table 2; Supplementary Material 5). For the whole-body transcriptome data, aminopeptidase N (APN) proteins were strongly enriched, with different genes within this family upregulated in each of the parental aphids (discussed further below). These trends were reinforced by comparison of parental transcriptomes in the heads RNA-Seq analyses where APN proteins were similarly enriched in both parents. The DE gene sets between the pooled VIR and AVR F1 samples indicated no enriched terms in the VIR data, and only a single term among the AVR upregulated genes: glucosidase II complex, that localises to the ER. These two gene sets are both relatively small (64 and 24 genes), reducing the likelihood of finding significant trends. Because very few significantly enriched terms were revealed by the initial GO analyses, we applied a lower stringency to inform wider trends in each of the DE gene sets. Here, we examined all terms for which at least two genes and a significant P value (<0.05) were returned. For the DE gene sets from RNA-Seq of heads, the majority of enriched terms were associated with the virulent N116 parent on both host genotypes. Although there was obvious redundancy of many terms, a substantial proportion (30-40%) for N116 relate to energy metabolism including mitochondria, TCA cycle, oxidative phosphorylation and lipid metabolism. In contrast, the PS01 enriched terms included several for protein processing including peptidases, proteolysis and protein glycosylation; and several for ATP-related transport (Supplementary Material 5). When each parental aphid genotype was compared separately for its differential responses to the two host genotypes (A17 and DZA), no significant terms were found for PS01, and only one weakly significant term for N116: polytene chromosome puffing. The equivalent GO analysis of whole body RNA-Seq data returned significantly enriched terms for both aphid genotypes, including several for protein modification (Supplementary Material 5).

**Table 2.** Significantly enriched GO terms within differentially expressed transcript and protein data. Terms enriched at FDR<0.05, after manual curation to remove redundancies, retaining the terms with lowest FDR. Full lists of enriched terms are in Supplementary Material 5.

| GO Category | Term | Ontology group | No. genes in DE set | P value | FDR |
| --- | --- | --- | --- | --- | --- |
| **Whole body RNA** |  |  |  |  |  |
| *N116 up*<br>0004177 | aminopeptidase activity | MF | 8 | 2.72E-06 | 0.0270 |
| *PS01 up*<br>0004177 | aminopeptidase activity | MF | 9 | 1.43E-07 | 0.0014 |
| 0017177 | glucosidase II complex | CC | 4 | 2.09E-06 | 0.0104 |
| *AVR F1 up*<br>0017177 | glucosidase II complex | CC | 3 | 2.42E-06 | 0.0240 |
| *VIR F1 up* | *No terms* |  |  |  |  |
| **Heads RNA** |  |  |  |  |  |
| *N116 up on A17*<br>0045271 | respiratory chain complex I | CC | 11 | 1.55E-07 | 9.63E-05 |
| 0005743 | mitochondrial inner membrane | CC | 22 | 2.01E-07 | 0.0001 |
| 0004177 | aminopeptidase activity | MF | 10 | 2.13E-06 | 0.0009 |
| 0016491 | oxidoreductase activity | MF | 31 | 3.24E-06 | 0.0012 |
| 0005875 | microtubule associated complex | CC | 25 | 1.72E-05 | 0.0052 |
| 0019395 | fatty acid oxidation | BP | 5 | 2.44E-05 | 0.0064 |
| 0042826 | histone deacetylase binding | MF | 4 | 0.00011 | 0.0275 |
| 0045239 | tricarboxylic acid cycle enzyme complex | CC | 3 | 0.00014 | 0.0327 |
| 0004448 | isocitrate dehydrogenase activity | MF | 3 | 0.00015 | 0.0338 |
| *N116 up on DZA*<br>0006635 | fatty acid beta-oxidation | BP | 5 | 1.05E-06 | 0.0082 |
| 0004177 | aminopeptidase activity | MF | 9 | 2.66E-06 | 0.0082 |
| 0004449 | isocitrate dehydrogenase (NAD+) activity | MF | 3 | 1.40E-05 | 0.0198 |
| 0006099 | tricarboxylic acid cycle | BP | 6 | 3.54E-05 | 0.0389 |
| *PS01 up on A17*<br>0004177 | aminopeptidase activity | MF | 10 | 7.77E-07 | 0.00771 |
| *PS01 up on DZA*<br>0004177 | aminopeptidase activity | MF | 11 | 5.59E-09 | 5.53E-05 |
| **Salivary gland proteins** |  |  |  |  |  |
| *N116 up*<br>0003983 | UTP:glucose-1-phosphate<br>uridylyltransferase activity | MF | 3 | 4.97E-06 | 0.0213 |
| *PS01 up* | *No terms* |  |  |  |  |

For the DE datasets from salivary gland proteomes, the lower stringency analysis revealed enrichment of distinct functional categories for each parental genotype. For N116, protein modification terms were prevalent including peptidase activity, serine-type endopeptidase inhibitor activity, negative regulation of protein metabolic process, aminopeptidase activity, protein kinase binding and regulation of protein phosphorylation. In contrast, for PS01, ATPase terms were predominant including several related to membrane transport, as also found in the PS01 heads RNA-Seq data (Supplementary Material 5).

### Exopeptidases are abundant in saliva, and the majority are DE between aphid genotypes

The saliva protein total and DE lists were much shorter, precluding formal GO analysis, but manual inspection indicated high proportions of exopeptidases: a total of 29 different proteins (Table 3), representing 22-34% of the protein list for each genotype. These were mainly APN proteins but also four members of the Angiotensin Converting Enzyme (ACE) family that are M2 metalloproteases with carboxypeptidase activity. The abundance of APNs in the saliva protein list broadly corroborates the major enriched GO categories detected in the transcriptome analyses.

**Table 3.**
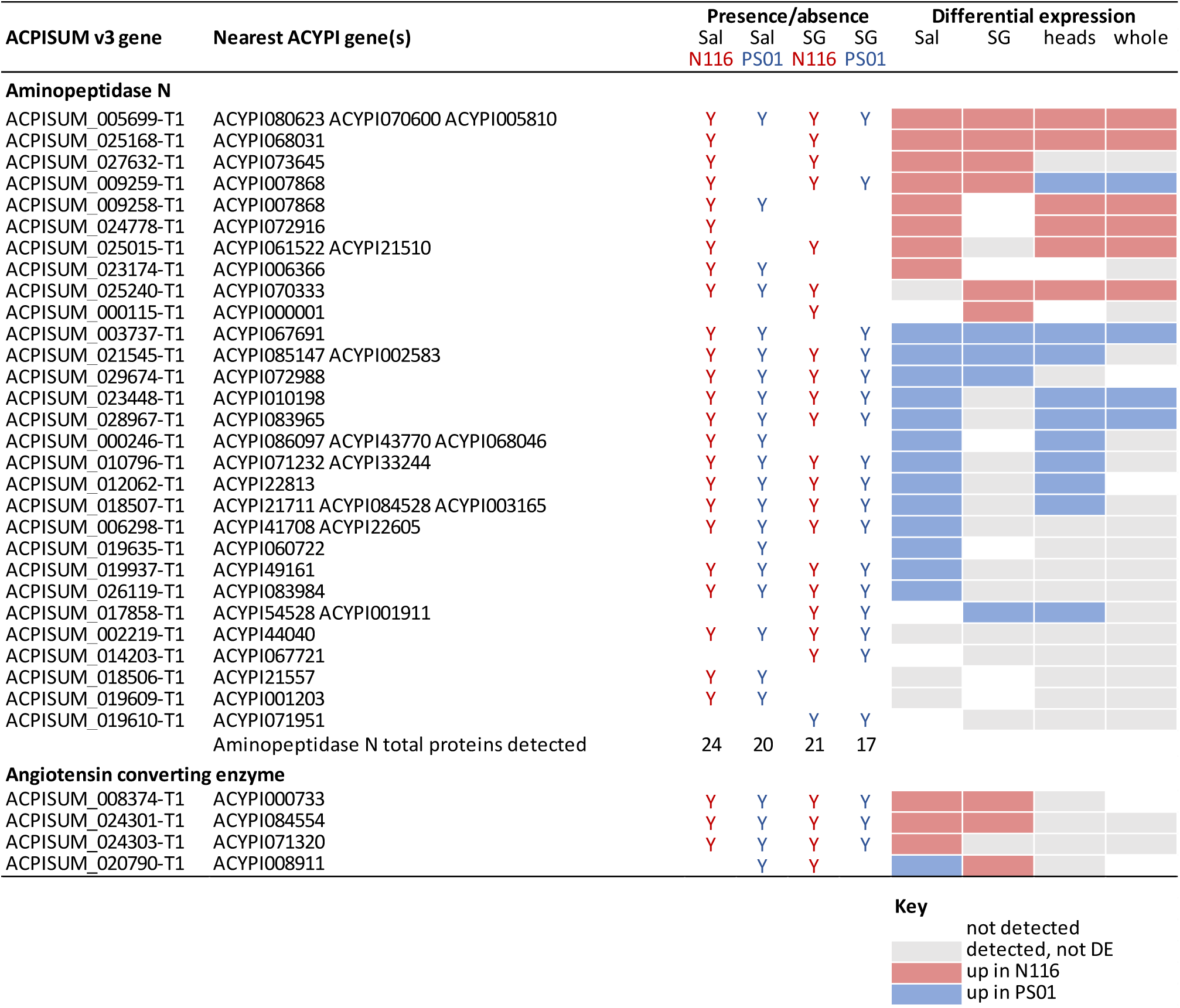
Comparison of expression patterns of exopeptidases detected in saliva and salivary glands. All detected proteins are listed, along with whether they were differentially expressed, and whether the patterns were also reflected in the transcriptomes. Sal = saliva; SG = salivary gland; Y = protein present.

Most of the exopeptidases detected from aphid saliva (23/29; 79%) were differentially abundant between the parental aphid genotypes. Twenty-two of the 29 saliva exopeptidases were also found in the salivary gland proteomes, with many showing the same direction of differential expression (9 APN, 2 ACE). Moreover, 15 (60%) of the APN proteins were DE in heads and/or whole body RNA-Seq samples (Table 3). Previous reports on pea aphid saliva and salivary gland components have also reported multiple APN and ACE proteins [12, 13, 32, 34]. Similar to our findings, one of these studies reported 11 APN genes that were differentially expressed in a biotype-specific manner, with five of these detected as proteins in saliva [13]. Taking all the evidence together, it is clear that the APN family is highly diversified in pea aphids and represents a major component of the salivary proteome by several measures: the high total number of proteins detected, many of these proteins are high abundance (13 of 20 top scoring in both N116 and PS01 saliva), and most are differentially expressed between aphid genotypes.

Aphid and mammalian ACE proteins have similar sequences and may have broadly similar functions as dipeptidases or by cleaving a single amino acid from the C terminus. However, mammalian ACE proteins are membrane anchored whereas aphid ACEs carry secretion signals, consistent with their detection in saliva. The exact catalytic functions and biological roles of aphid ACE and APN proteins remain to be determined. Cleavage of proteins and peptides could relate to targeting host proteins such as those involved in defensive sieve-tube blocking as shown at least for the atypical extrafascicular phloem exudate of cucurbits [35]. Alternatively, although there is currently no direct evidence, exopeptidases may act on other salivary protein components, for example to process effectors into active forms. Another non-mutually exclusive possibility is a role in aphid nutrition, with many insects using extra-organismal (extra-oral) digestion typical of arthropods including Hemiptera, enabling nutrition capture from large hosts/prey [32, 36]. Exopeptidases typically release N or C terminal single amino acids and dipeptides, potentially enabling supply of essential amino acids, some of which cannot be biosynthesised directly from the enzyme repertoires of hemimetabolous aphids.

### Multi-omic approaches to detecting candidate effectors

We compared the efficiencies of the four different experiments in terms of detecting aphid candidate effectors and related genes: RNA-Seq of heads and whole bodies, and proteomics of saliva and salivary glands. For all datasets, we focussed mainly on differential expression between the highly divergent parental clones N116 and PS01. Because saliva represents the “ground truth” of proteins predicted to be delivered into plant host tissues, we additionally considered saliva proteins that were detected but not DE. Although the proteomics methods are highly sensitive, there are likely to be some further low abundance salivary proteins that were not detected here. In addition, there may be some salivary proteins that are only expressed in response to aphids interacting with their host plants, and hence would not be found in artificial diet samples. Likewise, some proteins may not be stable under the artificial diet conditions. As a case study, we selected the significantly enriched exopeptidases, that comprised the large APN family and the smaller group of ACE proteins. We compared success of detecting genes from the saliva data in the other three experiments, and noted whether the same DE patterns were found (Table 3). The overall trends were broadly correlated, with 18/24 (75%) DE saliva proteins also found to be DE in at least one of the other approaches. Only two genes showed a mismatch in DE direction: ACPISUM_009259 between salivary gland and whole body; and ACPISUM_020790 between saliva and salivary gland. Individually, RNA-Seq of heads was the most effective experiment (14/24) at corroborating the DE saliva protein evidence, followed by RNA-Seq of whole bodies (10) and proteomics of salivary glands (8).

There are several reports where effectors are predicted from aphid salivary gland transcriptomes or proteomes, or other transcriptome datasets, typically filtering for presence of a signal peptide or other secretion motif, and absence of transmembrane domains [12–17]. For our exopeptidase data (Table 3), we detected an additional seven APNs in salivary gland proteomes or the transcriptome data, that were not found in saliva, of which five were DE in at least one dataset. Their absence from saliva indicates these proteins may be considered false positives for candidate effectors, although some low expressed proteins may go undetected. We considered which of the approaches was the most effective at detecting candidate effectors, and whether multiple omics approaches are advantageous, noting that all require substantial resource investment. Although saliva collection is an exacting and time-intensive procedure, saliva proteomics provided the greatest coverage of candidate effectors here, and quantitative analysis of mass spectrometry data enables robust assignment of differential expression. Of the other approaches, RNA-Seq of heads may be the most effective means to complement the saliva analyses by reinforcing evidence of differential expression, but in the work here did not greatly extend the effector lists per se.

## Conclusion

In this study, we demonstrated that transcriptomics and proteomics are both highly effective tools for discovering differentially expressed aphid genes and proteins. The protein subsets present in saliva are likely candidates for effectors with virulence and/or avirulence functions in host plants, and represent priorities for further study especially to determine if differential protein abundance is inherited into the segregating F1 aphid populations. Precise biochemical functions and host targets of most of these effectors are also currently unknown even in cases, such as the exopeptidases, where there are confident gene annotations. Exopeptidases are dominant in saliva by number of different proteins, by frequency of differential abundance, and by quantity. Likewise, there are many proteins of unknown function, with a substantial proportion found at high levels in saliva. Some of these unknown proteins may prove to be pivotal in explaining aphids’ unique and highly successful lifestyle as phloem feeders.

## Methods

### Aphids and crossing

Pea aphid (*Acyrthosiphon pisum*) clones were maintained on tic bean (*Vicia faba minor*) as described in [26]. Parental genotypes were PS01 and N116. PS01 is a biotype adapted to *Pisum sativum* whereas N116 is adapted to *Medicago sativa* [26]. Reciprocal crosses were made between PS01 and N116 to generate F1 hybrid populations, following the protocol of [27]. In brief, parthenogenetic females were induced to generate sexual forms by transfer to short days and lower temperatures to simulate autumn. Eggs resulting from controlled matings were collected onto moist filter paper in petri dishes, and subjected to 90 to 105 days at 4°C to induce exit from diapause. Individual hatchlings were subsequently used to generate multiple parallel clonal F1 lineages. Parents and progeny were genotyped with a set of seven microsatellite markers [22] to verify correctness of crosses. All new F1 progeny were maintained for at least three generations before testing performance on different host plants.

### Plants and assessment of virulence

Based on previous findings [27], PS01 aphids are avirulent on *Medicago truncatula* J A17 that carries the resistance QTL, *RAP1* [28]. Near isogenic lines (NILs) derived from a cross (LR4 [29]) between A17 and *M. truncatula* DZA315.16 were also used. PS01 is likewise incompatible with the resistant NIL (RNIL), but is compatible with the susceptible NIL (SNIL) and with DZA315.16. N116 aphids are compatible with all these genotypes. F1 progeny were tested for virulence on both A17 and RNIL, based on [26]. Briefly, five nymphs of each clone were infested onto ten A17 or RNIL plants, then scored for survival and production of new nymphs 10 d later. At least 40 F1 clones each of PS01 x N116 and N116 x PS01 were screened. An overall virulence index was adapted from a calculation proposed in [37]:

Virulence index = log2 (mean number surviving out of 5 x number of nymphs produced + 1)

Virulent (VIR) clones were defined as index >4 and >5 on A17 and RNIL, respectively, and avirulent (AVR) clones were correspondingly defined as index <2 and <4. The different category thresholds on A17 and RNIL reflect the latter’s slightly lower resistance. Clones falling into the same phenotype category (VIR or AVR) on both A17 and RNIL were then subject to a further confirmation screen where survival on A17 and RNIL was counted 5 d after infestation. In the confirmation experiment, four plants were used for each aphid x host combination, with five aphids infested onto each plant. Cutoffs were >80% survival for virulence on both hosts, and <20% and <70% for avirulence on A17 and RNIL, respectively. A few F1 clones showed relatively high survival at 5 days but had very weak growth, and therefore were categorised as AVR. Only F1 clones displaying the same phenotype category on all screening experiments were used subsequently in molecular experiments.

### Sampling for RNA-Seq

Heads experiment: Young adult aphids of clones N116 and PS01, cultured on *Vicia faba minor*, were infested onto either A17 or DZA315.16 *M. truncatula* plants for 24 h, then heads (40 per sample) were dissected and frozen immediately on dry ice then stored at -80°C. Three replicates were done for each aphid x plant combination.

Whole body experiment: Samples were parental aphid clones (N116 and PS01) and pools of VIR and AVR F1 progeny. Aphids of each individual genotype, age 2 to 3 d, were placed on independent A17 plants for 24 h then frozen in liquid nitrogen and stored at -80°C until processing. A total of 22 VIR and 22 AVR F1 aphid clones were collected individually, before pooling five aphids of each genotype to comprise one sample. Five biological replicates were analysed for both parental and pooled F1 genotypes.

### RNA extraction, library preparation and sequencing

Heads were dissected and processed as described in [16]. Total RNA was extracted using a plant RNA extraction kit (Sigma-Aldrich). Illumina TruSeq stranded mRNA-Seq libraries were sequenced at the Genome Sequencing Unit at the University of Dundee on an Illumina HiSeq 2000.

RNA for the BSA-RNA-Seq analysis was isolated from three two to three day old nymphs of parental lines (N116, PS01), 22 VIR F1 lines and 22 AVR F1 lines, using the Norgen Plant and Fungal RNA kit (Sigma E4913). The RNA isolation followed the instructions of the company supplementing Lysis buffer C with ß-mercaptoethanol. An on-column DNase digest was performed (RNase-Free DNase Set, Qiagen) and the concentration of each sample determined via a Qubit fluorometer with the QubitTM RNA Broadrange (BR) assay kit (Thermo Fisher Scientific). Samples corresponding to five replicates of each of the parental lines and the VIR and AVR F1 pools were used to generate a total of 20 Illumina TruSeq stranded mRNA-Seq libraries which were sequenced in 150 bp paired-end mode on an Illumina HiSeq4000 at Edinburgh Genomics.

### RNA-Seq data processing and visualisation

Illumina RNA sequence reads were subjected to quality control using FastQC. The reads were the trimmed using Trimmomatic (version 0.32) Q15, min length 55. The trimmed fastq files were then quasi mapped to the nucleotide gene sequences for the pea aphid using salmon version 1.1. For the pilot study, STAR (2.4.1b) [38] was used to map the reads to the pea aphid genome and HTseq counts was used to quantify the gene expression using AphidBase_OGS2.1b gene annotations.

Clone-specific *de novo* RNA-Seq assemblies (from both the heads and whole-body studies) were individually and collectively generated using Trinity version 2.9.1. All the data were pooled into one for the “collective” assembly, which was used for transcript differential expression analysis. The individual assemblies were used for gene prediction at a later stage. All RNA-Seq assemblies were quality filtered using Transrate to reduce the probability of mis-assembled transcripts. Predicted coding sequences were generated using TransDecoder (with PFAM and BLAST guides). Diamond was used to search against GenbankNR database. Differential expression analysis was performed using EdgeR. Heatmaps and expression profile clustering were generated using the ptr script from within the Trinity package.

During early analysis, following visual assessment of RNA-seq read mapping and initial differential expression results, we found that the original pea aphid gene predictions (AphidBase_OGS2.1b) and the gene predictions from [39] did not fully match those generated by the de novo transcriptome assemblies. Therefore, gene annotation was re-predicted on the published pea aphid genome (OGS2.1b) to improve the accuracy of the gene models. Funannotate, in Other Eukaryotic mode, was used to predict the genes using the *de novo* RNA-Seq assembly generated above, with RNA-Seq data mapped using STAR (see above). A total of 29,930 genes were assigned codes in the format ACPISUM_0xxxxx, with the annotations provided at doi.org/10.5281/zenodo.11103500 [40].

To assign the various gene calls from the original genome assembly, bedtools intercept was used to identify genes with overlapping coordinates. If the genes overlapped, then they were considered the same gene. A simple BLAST approach could not be used here due to the duplicated nature of aphid assemblies. A combination of reciprocal best BLAST hit, Orthofinder and MCL clustering were used to assign genes between the clones as orthologues.

### Saliva Collection

For proteomics samples, N116 and PS01 were maintained separately on *Vicia faba* c.v. The Sutton, grown in standard potting compost and kept at 20°C and a photoperiod of 16-h light/8-h dark. Approximately 3,000 mixed aged aphids were positioned on 30 perspex rings (radius 4.5 cm, height 5 cm), each containing 4.5 ml of a chemically-defined diet, formulation A from [41], held between two stretched sheets of Parafilm^TM^. The aphids were reared on the diets at 20°C with 18h light and 6h dark for 24 h after which the diets were pooled and collected and stored at -80°C until required. Four independent replicates were produced by pooling the collected diet from two daily collections (approximately 150 ml). Pooled diets were concentrated using a Vivacell 250 Pressure Concentrator (Sartorius Mechatronics, UK) using a 5000 Da molecular weight cut-off (MWCO) polyethersulfone (PES) membrane. When the final volume had reached 5 ml it was removed and 1 ml of filtered sterilised PBS (phosphate-buffered saline) supplemented with Roche cOmplete^TM^ protease inhibitor cocktail (PIC) was added. The resulting mixture was further concentrated to approximately 250 μl using a Vivaspin 6 centrifuge concentrator (Sartorius Mechatronics, UK) with a 5000 Da MWCO PES membrane, purified using a 2D Clean-up Kit (GE HealthCare) following the manufacturer’s instructions. The resulting protein pellet was suspended in 25 μl 6 M urea, 2 M thiourea, 0.1 M Tris-HCl, pH 8.0 and re-quantified using the Qubit Fluorometer. Four independent biological replicates per genotype were subjected to mass spectrometry.

### Salivary glands

The salivary glands from 14-16 day old adult aphids of N116 and PS01 were dissected in ice-cold PBS and transferred to 60 µl PBS supplemented with PIC. Forty pairs of salivary glands were pooled per replicate and homogenized with a motorised, disposable pestle. Sixty microliters of 12 M urea, 4 M thiourea, and PIC was added and the samples were homogenised further and centrifuged at 9,000 × *g* for 5 min to pellet cellular debris. The supernatant was removed and quantified, and 100 µg of protein was purified using a 2D Clean-up Kit (GE HealthCare) following the manufacturer’s instructions with the exception that 400 μl of precipitant and co-precipitant were used in the first step. The resulting protein pellet was re-suspended in 30 μl 6 M urea, 2 M thiourea, 0.1 M Tris-HCl, pH 8.0 and re-quantified using the Qubit Fluorometer. Four biological replicates per genotype were subjected to mass spectrometry.

### Protein sample digestion for mass spectrometry

The digestion protocol was the same for both saliva and salivary gland samples and involved the addition of 50 μl ammonium bicarbonate, reduction with 0.5 M dithiothreitol at 56°C for 20 min and alkylation with 0.55 M iodoacetamide at room temperature for 15 min, in the dark. One μl of a 1% w/v solution of ProteaseMax Surfactant Trypsin Enhancer (Promega) and 1 μg of Sequence Grade Trypsin (Promega) were added, then samples were incubated at 37°C for 18 h. Digestion was terminated by adding 1 μl of 100% trichloroacetic acid (Sigma Aldrich) and incubating at room temperature for 5 min. Samples were centrifuged for 10 min at 13,000 *x g* and the supernatant was removed to new microcentrifuge tubes.

### Mass spectrometry and proteomic data analysis

One μg of digested peptide was loaded onto a Dionex Ultimate 3000 (RSLCnano) chromatography system connected to a QExactive (ThermoFisher Scientific) high-resolution accurate mass spectrometer. Peptides were separated by an increasing acetonitrile gradient on a Biobasic C18 PicofritTM column (100 mm length, 75 µm ID), using 120 and 50 min reverse phase gradients for salivary glands and saliva, respectively, at a flow rate of 250 nl min^-1^. All data were acquired with the mass spectrometer operating in automatic data dependent switching mode. A high-resolution MS scan (300-2000 Da) was performed using the Orbitrap to select the 15 most intense ions prior to MS/MS.

Protein identification and normalisation was conducted using the Andromeda search engine in MaxQuant (version 1.6.17.0; http://maxquant.org/) to correlate the data against the predicted protein set generated in this study (ACPISUM_Proteins; 30891 entries) using default search parameters for Orbitrap data. False Discovery Rates were set to 1% for both peptides and proteins and the FDR was estimated following searches against a target-decoy database. Two searches were conducted for both N116 and PS01 saliva and salivary glands. The first involved a combined search of the raw files for each genotype separately to generate comprehensive proteomes for the saliva or salivary gland (hereafter All Identified Proteins). The second involved a quantitative search of the raw files for all biological replicates (n=4) for the saliva or salivary glands. Quantitative and statistical analyses were conducted in Perseus (Version 1.6.1.1 http://maxquant.org/) using the normalized label-free quantitation (LFQ) intensity values from each sample. The data were filtered to remove contaminants, and peptides identified by site. LFQ intensity values were log_2_ transformed and samples were allocated to their corresponding groups. A data imputation step was conducted to replace missing values with values that simulate signals of low abundant proteins chosen randomly from a distribution specified by a downshift of 2.1 times the mean standard deviation (SD) of all measured values and a width of 0.1 times this SD. Normalized intensity values were used for principal components analysis. A two-sample t-test was performed using a cut-off value of p ≤ 0.05 to identify statistically significant differentially abundant (SSDA) proteins. Volcano plots were produced by plotting –Log p-values on the y-axis and Log_2_ fold-change values on the x-axis to visualize differences in protein abundance between the two genotypes.

### Gene annotations and Gene Ontology analysis

Secretion signal properties were predicted using SignalP4.1 [42]. Non-annotated genes were defined as those with the following descriptors: hypothetical protein, uncharacterized protein, NA or ACYPIxxxxx without any other assigned function. GO enrichment analyses were performed using GOseq [43].

### Data availability

Genome annotations: zenodo.org/records/11103500 [40]

RNA-Seq: Pea aphid clones N116 and PS01 reared on *Medicago truncatula* A17 and DZA315.16, dissected heads: BioProject PRJNA757589, ncbi.nlm.nih.gov/bioproject/PRJNA757589/

RNA-Seq: Pea aphid clones N116, PS01 and bulk F1 hybrid progeny reared on *Medicago truncatula*

A17, whole body samples: BioProject PRJNA757896, ncbi.nlm.nih.gov/bioproject/PRJNA757896

Scripts: github.com/peterthorpe5/Pea_aphid_on_medicago_DZA_A17

Proteomics: mass spectrometry data have been deposited to the ProteomeXchange Consortium via the PRIDE partner repository [44], dataset identifiers PXD053355 and PXD053620.

## Author contributions

JB, CT, JC, PT, SA and RLC designed the experiments. PT, SA, RLC, ND, JCS, JI and SK conducted the experiments. PT, SA, RLC, JB, CT, JC, ND and JCS analysed the data. CT, JB, JC and PT wrote the paper. All authors approved the submitted manuscript.

## Supplementary Materials

Supplementary Material 1. F1 aphid phenotyping.

Supplementary Material 2. Read mapping summary.

Supplementary Material 3. RNA-Seq Head and whole body differentially expressed genes.

Supplementary Material 4. Salivary gland and saliva proteomics.

Supplementary Material 5. GO enrichment

