## Supplementary Material 2 for "Multi-omics approaches define novel aphid effector candidates associated with virulence and avirulence phenotypes"

### Slide 1
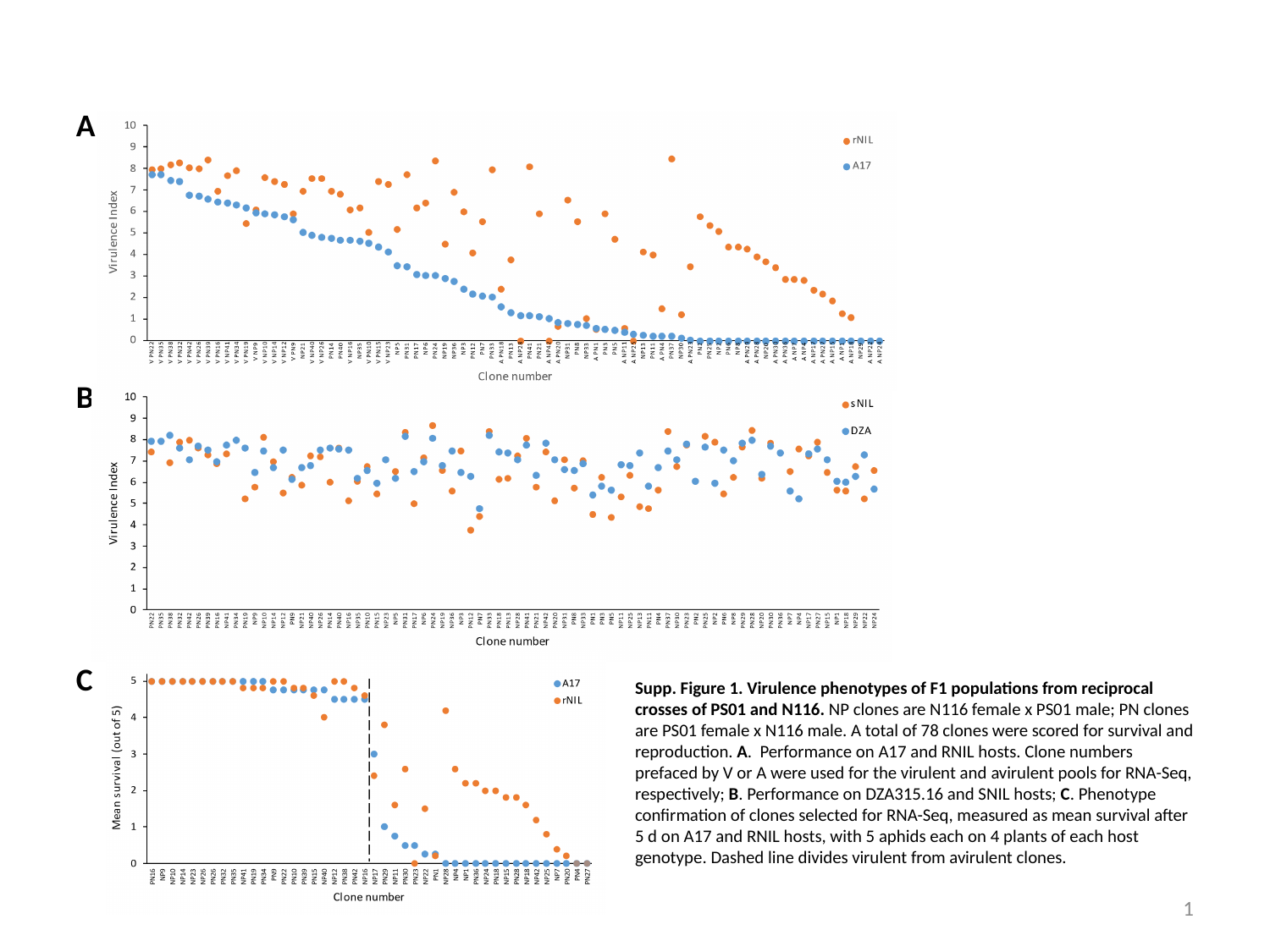

A
B
C
Supp. Figure 1. Virulence phenotypes of F1 populations from reciprocal crosses of PS01 and N116. NP clones are N116 female x PS01 male; PN clones are PS01 female x N116 male. A total of 78 clones were scored for survival and reproduction. A. Performance on A17 and RNIL hosts. Clone numbers prefaced by V or A were used for the virulent and avirulent pools for RNA-Seq, respectively; B. Performance on DZA315.16 and SNIL hosts; C. Phenotype confirmation of clones selected for RNA-Seq, measured as mean survival after 5 d on A17 and RNIL hosts, with 5 aphids each on 4 plants of each host genotype. Dashed line divides virulent from avirulent clones.
1
