## Supplementary Material 3 for "Multi-omics approaches define novel aphid effector candidates associated with virulence and avirulence phenotypes"

**Supplementary Material 2. Summary of RNA-Seq read mapping**

A. **Aphid heads experiment**

| **Sample** | **Number of raw read pairs** | **Uniquely mapped reads** | **Uniquely mapped reads %** | **% of reads unmapped** |
| --- | --- | --- | --- | --- |
| N116_A17_1 | 13207408 | 9601671 | 88.28% | 0.23% |
| N116_A17_2 | 12407873 | 9013436 | 88.98% | 0.21% |
| N116_A17_3 | 11925546 | 8695377 | 88.53% | 0.14% |
| N116_DZA_1 | 11873230 | 8637078 | 90.87% | 0.12% |
| N116_DZA_2 | 14274392 | 10278702 | 87.85% | 0.18% |
| N116_DZA_3 | 13861349 | 9968169 | 87.57% | 0.19% |
| PS01_A17_1 | 14744045 | 10620809 | 88.32% | 0.23% |
| PS01_A17_2 | 13151054 | 9334734 | 87.10% | 0.33% |
| PS01_A17_3 | 13949759 | 9624856 | 87.25% | 0.26% |
| PS01_DZA_1 | 14139437 | 10154324 | 87.78% | 0.27% |
| PS01_DZA_2 | 9641233 | 6835657 | 87.74% | 0.30% |
| PS01_DZA_3 | 12716143 | 9028027 | 87.58% | 0.27% |

B. **Whole aphid experiment**

| **Sample** | **Number of reads** | **Uniquely mapped reads** | **Uniquely mapped reads %** | **% of reads unmapped** |
| --- | --- | --- | --- | --- |
| avr-rep1 | 18926678 | 16346937 | 86.37% | 7.38% |
| avr-rep2 | 23020445 | 19841290 | 86.19% | 8.09% |
| avr-rep3 | 26800376 | 21904963 | 81.73% | 8.58% |
| avr-rep4 | 22119898 | 19174155 | 86.68% | 7.81% |
| avr-rep5 | 24411325 | 21045037 | 86.21% | 7.60% |
| N116-rep1 | 23671278 | 18311195 | 77.36% | 16.58% |
| N116-rep2 | 18671032 | 14546128 | 77.91% | 17.16% |
| N116-rep3 | 22285687 | 17316495 | 77.70% | 17.81% |
| N116-rep4 | 26015956 | 21001704 | 80.73% | 16.29% |
| N116-rep5 | 19794870 | 15594144 | 78.78% | 15.62% |
| PS01-rep1 | 20158864 | 18068560 | 89.63% | 4.10% |
| PS01-rep2 | 18435849 | 15942153 | 86.47% | 4.38% |
| PS01-rep3 | 21324177 | 19277973 | 90.40% | 4.33% |
| PS01-rep4 | 20609372 | 18018966 | 87.43% | 4.01% |
| PS01-rep5 | 18901306 | 17272176 | 91.38% | 3.87% |
| vir-rep1 | 23269113 | 20377850 | 87.57% | 7.02% |
| vir-rep2 | 19590868 | 17006965 | 86.81% | 7.29% |
| vir-rep3 | 18974382 | 15718127 | 82.84% | 7.46% |
| vir-rep4 | 19057070 | 16558858 | 86.89% | 7.73% |
| vir-rep5 | 19449710 | 16622072 | 85.46% | 6.98% |
